# Development of a genetically encoded melanocortin sensor for high sensitivity intravital imaging

**DOI:** 10.1101/2025.08.08.669345

**Authors:** Yoon Namkung, Tal Slutzki, Joao Pedroso, Xiaohong Liu, Paul V. Sabatini, Maia V. Kokoeva, Stéphane A. Laporte

**Affiliations:** Department of Medicine, Division of Endocrinology and Metabolism, Research Institute of the McGill University Health Center, McGill University, Montréal, QC H4A 3J1, Canada; Integrated Program in Neuroscience, McGill University, Montréal, QC H3A 2B4, Canada; Department of Pharmacology and Therapeutics, McGill University, Montréal, QC H3G 1Y6, Canada

**Keywords:** α-MSH, Melanocortin receptors MC1 and MC4, Circularly permuted green fluorescent protein, Intraviral brain imaging sensor, Central appetite regulation, Hypothalamus

## Abstract

The central melanocortin system, composed of peptides derived from pro-opiomelanocortin (POMC) such as the melanocyte-stimulating hormones (α-, β-, γ-MSH) and melanocortin 4 receptors (MC4R), along with the agouti-related protein (AgRP), plays a pivotal role in controlling energy balance. To elucidate the dynamic role of α-MSH release in regulating appetite, specific, sensitive, and spatiotemporally resolved genetic sensors are required. The melanocortin 1 receptor (MC1R) scaffold was leveraged for its robust plasma membrane expression, high affinity for melanocortin peptides and low affinity for AgRP to design a α-MSH selective sensor for *in vivo* use. This was achieved by integrating circularly permuted green fluorescent protein (cpGFP) into the receptor, which we named **<u>Fl</u>**uorescence **<u>A</u>**mplified **<u>Re</u>**ceptor sensor for Melanocortin (FLARE_MC_). The FLARE_MC_ sensor shows high potency and selectivity in heterologous and homologous expressing cells for α-MSH and the synthetic melanocortin agonist MTII but not to the inverse agonist AgRP. The sensor exhibited impaired signaling, with reduced G protein activation, no β-arrestin coupling, and failed to internalize upon agonist stimulation. *In vitro*, FLARE_MC_ displayed high photostability and reversible photoactivation. These properties suggest that the FLARE_MC_ is suitable for long-term activity recording in the brain without desensitizing or interfering with endogenous melanocortin receptor signaling. When expressed in the paraventricular nucleus (PVN) of the mouse hypothalamus, the primary site of anorexigenic α-MSH signaling, FLARE_MC_ demonstrated its effectiveness in detecting changes associated with melanocortin responses *in vivo*. FLARE_MC_ enables the study of melanocortin system in cultured cells and intravitally. This first of its class highly sensitive melanocortin sensor will serve as a valuable tool to advance our understanding of the complex dynamics governing melanocortin-dependent appetite regulation and related processes in the brain.

## Introduction

The melanocortin system is a critical regulator of organismal energy homeostasis. Central to this system are the anorexigenic pro-opiomelanocortin (POMC) and orexigenic agouti-related protein (AgRP)-expressing neurons of the arcuate nucleus, which signal onto melanocortin 4 receptor (MC4R)-expressing neurons located in paraventricular nucleus (PVN) of the hypothalamus (1, 2). MC4R is a G protein-coupled receptor (GPCR) belonging to the melanocortin receptor family, which comprises five members (MC1R–MC5R). Like other melanocortin receptors, MC4R signals primarily through Gas and is activated by POMC-derived peptides— including adrenocorticotropic hormone (ACTH) and a-, B-, and y-melanocyte–stimulating hormones (MSH). Its activity is antagonized by the endogenous inverse agonist AgRP (3). Importantly, alterations of the melanocortin system have similar phenotypic outcomes in both mice and humans. Accordingly, loss of a-MSH production due to *e.g.* POMC gene disruption or deficiency in MC4R cause severe obesity in mouse models and humans (4–6).

Owing to their crucial role in regulating energy balance, AgRP and POMC neurons are highly sensitive to changes in nutritional status and food availability. AgRP neuron activity increases with fasting and reduces following the sensory detection of nutrients within the gastrointestinal tract or even when animals are visually exposed to feeding cues (7), whereas POMC neuron activity increases following sensory detection of food (8, 9). While these studies show that nutrient sensing rapidly modulates POMC and AgRP neuron activity, how this translates into changes in melanocortin hormone release and downstream receptor signaling remains poorly understood.

To monitor *in vivo* changes in melanocortin release, particularly a-MSH, with high temporal resolution, we developed a sensor based on MC1R, which contrary to MC4R, offers robust plasma membrane expression, high affinity for melanocortin, and low affinity for AgRP (3). We named this sensor **<u>Fl</u>**uorescence **<u>A</u>**mplified **<u>Re</u>**ceptor for Melanocortin (FLARE_MC_) and show that FLARE_MC_ is highly sensitive and selective to the POMC derivative a-MSH, with low responsiveness to AgRP. Additionally, expressing the FLARE_MC_ in the PVN of mice efficiently detects pharmacologic changes in melanocortin. This sensor represents a useful tool to further investigate the melanocortin system and its regulation of energy balance.

## Results

### MC1R is a suitable scaffold for the development of a sensitive genetically encoded melanocortin sensor

To develop a genetically encoded melanocortin sensor for *in vivo* usage, we initially planned to introduce a circular permutated GFP (cpGFP) into the third intracellular loop (ICL3) of the mouse MC4R (mMC4R). Similar strategies were previously employed to generate genetically encoded sensors for different neurotransmitters and ligands acting on GPCRs (10–19). These sensors are based on the principal that conformational changes within the receptor following ligand binding alters the arrangement of the integrated cpGFP, resulting in stabilization of the chromophore and changes in fluorescence (Figure 1A). We first characterized the distribution of the mMC4R in cells and its tolerance to introducing a GFP in its sequence by adding the fluorescent protein into a region of the receptor believed to be least disruptive, such as the C-terminus, and comparing it to the mMC1R with similar fusion of GFP (mMC1R-EGFP). We observed mMC4R-EGFP was predominantly intracellularly localized when expressed in HEK293 cells (Figure 1B) at basal state, which contrasted the robust plasma membrane (PM) localization of mMC1R-EGFP. Distribution of the MC4R, including its intracellular localization at basal state has been reported previously (20, 21). We reasoned that MC4R’s predominant intracellular localisation would impede effectiveness in detecting a-MSH *in vivo*, because of insufficient receptor density at the PM of neurons. Indeed, we observed intracellular accumulation with many cpGFP-mMC4R constructs (Data not shown). Moreover, incorporating a cpGFP within the ICL3 region of mMC4R may critically diminish its ligand affinity because of G proteins uncoupling, mirroring observations seen with other genetically engineered cpGFP-GPCR sensors (12, 15). We therefore shifted our effort to developing a mMC1R-based sensor, which exhibited predominant localisation on the PM (Figure 1B) and displays the highest intrinsic affinity for a-MSH among the melanocortin receptors, including MC4R (3). Importantly, since cpGFP-GPCR sensors generally have reduced signaling capabilities (11, 12, 15), a cpGFP-mMC1R sensor is not expected to disrupt signaling of MC4R in cells where it is natively expressed, when acting as sensitive proxy for a-MSH detection *in vivo*. Because MC1R has a short ICL3 similar to the CB1 receptor, we opted for a similar strategy and inserted the cassette of the GPCR-Activation-Based (GRAB) CB1 receptor sensor (eCB2.0) (11), which includes the cpGFP flanked by linkers between residues Q221^5.74^ and G232^6.28^, into mMC1R. This corresponds to positions R311^5.75^ and R336^6.28^ in the CB1 receptor (Ballesteros-Weinstein numbering in upper case, (22)) (Figure 1C). Moreover, the Q221^5.74^ in mMC1R was substituted for an Arg to mimic the hMC1R residue position and the last residue before the linker in the GRAB_eCB2.0_ cassette. We also tried to replace the ICL3 of the mMC1R with that of different GRAB-cpGFP sensors such as the Dopamine D2 (GRAB_DA2m_) (16) or the Norepinephrine (GRAB_NE1m_) (12) receptors, which contain the cpGFP flanked by the respective ICL3’s receptor sequence (Figure 1D). GRAB_eCB2.0_ cassette insertion in mMC1R generated a sensor that was efficiently expressed at the PM and produced both a good basal signal for receptor detection and the best signal-to-noise fluorescence (ΔF/F_0_) ratio when stimulated with the a-MSH analogue MTII (Figure 1D); hence we selected this version for further optimization.

**Figure 1:**
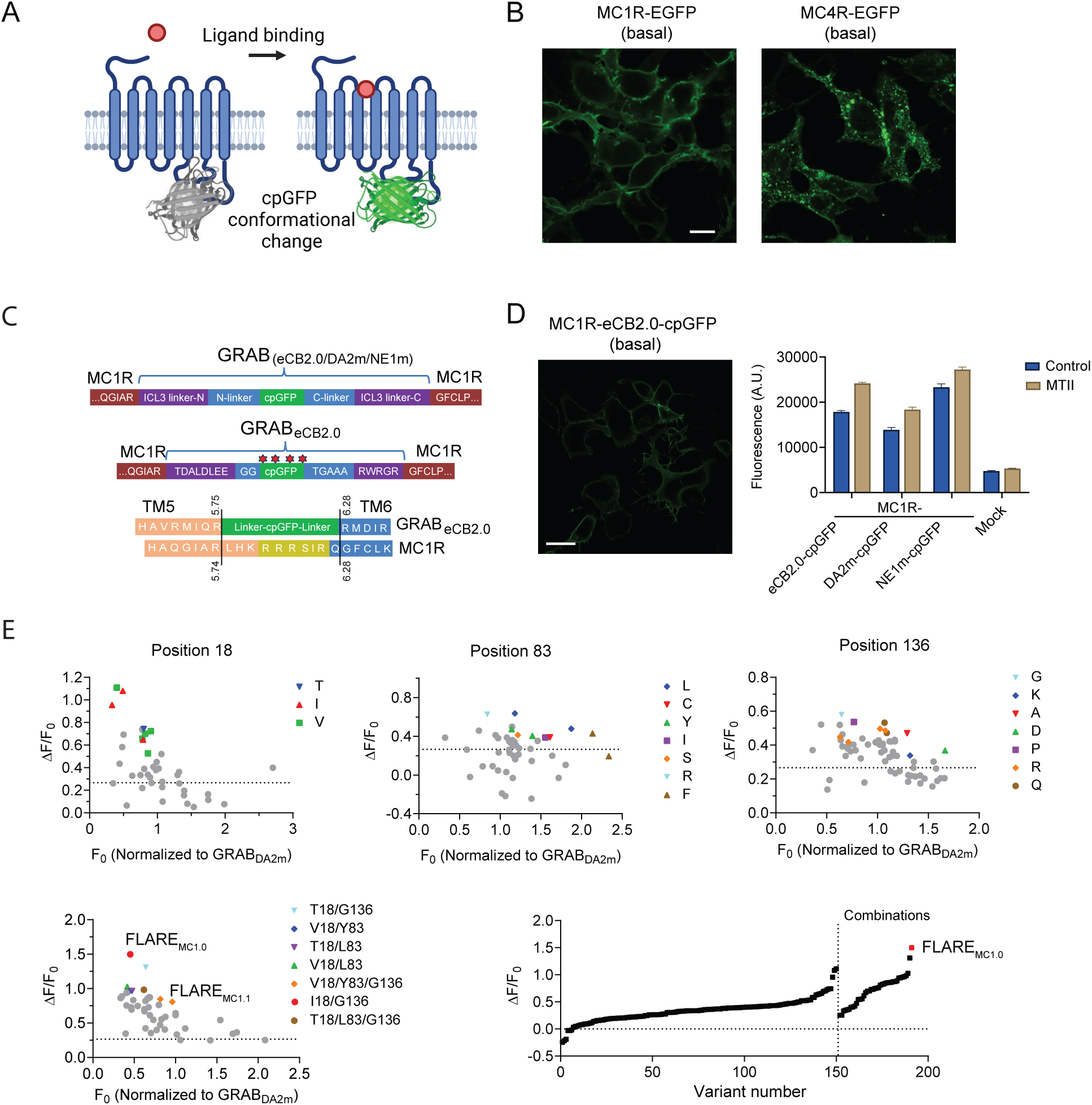
**Generation and optimization of the FLARE_MC_ sensors**. (**A**) Schematic diagram of circular permutated enhanced green fluorescent proteins (cpGFP)-based GPCR sensors, including the FLARE_MC_. Image was created with www.biorender.com. (**B**) Cellular localization of MC1R and MC4R receptor tagged with EGFP in their C-terminus, at basal state. HEK293 cells were transfected with either MC1R-EGFP or MC4R-EGFP constructs, and confocal microscopy was conducted 48 hours post-transfection. (**C**) Alignment of eCB2.0 sensor cpGFP junction with ICL3 junction of mMC1R. Vertical lines indicate where cpGFP and linker were introduced. (**D**) Confocal image of MC1R containing GRAB_eCB2.0_’s cpGFP with flanking regions including ICL3 (*left*). Cells were transfected with MC1R with its ICL3 replaced with cpGFP with flanking regions of either eCB2.0, DA2m, or NE1m sensor. Shown are total fluorescence in cells at basal (control) and with 10 μM MTII treatment (*right*). (**E**) Saturation-mutagenesis with amino acid substitutions at position 18, 83, and 136 of cpGFP. F_0_ values were normalized to those of the GRAB_DA2m_ sensor. Shown are the screened mutants, including FLARE_MC1.0_ and FLARE_MC1.1_, their responses at each position, and their overall ranking based on increased fluorescence. Scale bars (**B** and **D**), 10 μm.

### Residue-specific mutagenesis of cpGFP enhances sensor performance

We next aimed to improve the ligand-induced, conformationally-driven fluorescence of the sensor by targeting the cpGFP moiety. Critical residues were chosen from the cpGFP alignment which showed the greatest variability between different cpGFP-based sensors (13) (Figure 1C and Figure S1). Positions 18, 83, 136, and 208 were targeted for site-saturation mutagenesis, with random alterations conducted at each designated site (Figure S1) and to diversify the sequence, position 136 mutagenesis was done with cpGFPs from either GRAB_eCB2.0_ or GRAB_DA2m_ as templates. Between 48 and 65 clones for each position substitution at the four sites in cpGFP were screened, covering a widespread range of all 19 possible amino acid substitutions. Variants demonstrating optimal fluorescence dynamics at each site were identified and subsequently revalidated for their fluorescent changes. Substituting position 18 in cpGFP with Thr, Ile, or Val, produced the highest ΔF/F_0_ florescence increase (Figure 1E), while the best fluorescence increase occurred with Leu substitutions at position 83 and Gly substitutions at position 136, respectively. Position 208 exhibited low tolerance to substitutions, with approximately 60% of variants showing reduced fluorescence upon introduction of different residues at this site (*e.g.*, 20 out of 34 variants were inactive). That position was therefore not pursued for optimization. We next combined different amino acid substitutions at the first 3 positions of the cpGFP in mMC1R (Figure 1E). The Ile^18^/Gly^136^ substitutions, which we named FLARE_MC1.0_ showed the lowest basal fluorescence as compared to GRAB_DA2m_ and produced the most profound agonist-mediated changes in the receptor fluorescence with an increase of ∼150% (Figure 1E). Other clones such as FLARE_MC1.1_ (V^18^/Y^83^/G^136^ substitution) had similar basal fluorescence as compared to GRAB_DA2m_ but produced a reduced signal as compared to FLARE_MC1.0_. The dynamic range of the FLARE_MC1.0_ sensor was similar to those of other GRAB sensors, which were previously established for *in vivo* studies (12, 14, 15). We therefore proceeded to further characterize this sensor *in vitro*, hereafter referring to it as FLARE_MC1.0_, or simply FLARE_MC_ (Figure S2).

### FLARE_MC_ enables *a*-MSH and MTII detection independently of the sensor signaling

We initially characterized FLARE_MC1.0_ pharmacologically and functionally in heterologous cells, evaluating its ligand responsiveness, including to melanocortin, and its coupling to canonical G protein pathways. MTII or a-MSH increased fluorescence of FLARE_MC1.0_ at the PM in HEK 293 cells (Figure 2A). Dose-response curves revealed half-maximal effective concentrations (EC_50_) for FLARE_MC1.0_ sensors of approximately 12.5 nM and 34.8 nM, respectively (Figure 2B), consistent with the known high affinities and potencies of a-MSH and MTII for the MC1R (3). Moreover, FLARE_MC1.1_ exhibited a similar pattern of relative sensor activity in response to a-MSH and MTII as FLARE_MC1.0_, albeit with reduced amplitude (e.g. Emax), suggesting that the observed difference in agonist potency is unlikely due to clonal selection (Figure 2B). To evaluate the signaling capacity of the FLARE_MC_ sensors relative to mMC1R and mMC4R, we measured Gs-dependent cAMP production using an EPAC-based Bioluminescence Resonance Energy Transfer (BRET) sensor (23) (Figure 2C). Both MTII and a-MSH potently promoted cAMP production following challenges of cells expressing mMC1R or mMC4R with EC_50_ of 1.6 nM and 4.8 nM, and 4.7 nM and 11.5 nM, respectively. However, maximal response of the FLARE_MC1.0_ promoted by MTII or a-MSH was reduced by more than 75% with a substantial decrease in potency of 60-and 35-fold (EC_50_ of 97.5 nM and 166.7 nM for MTII or a-MSH, respectively) as compared to MC1R. FLARE_MC1.1_ showed similar reduction in potencies for MTII-mediated cAMP as compared to FLARE_MC1.0._ However, signaling responses reached 50% and 75% of those observed with mMC1R when activated by MTII and a-MSH, respectively (Figure 2C); suggesting FLARE_MC1.1_ retains signaling capacity. We prioritized FLARE_MC1.0_ for further characterization due to its superior dynamic range (Figure 2B) and lower intrinsic G protein activity (Figure 2C), which would minimize interference with endogenous melanocortin receptor signaling.

**Figure 2:**
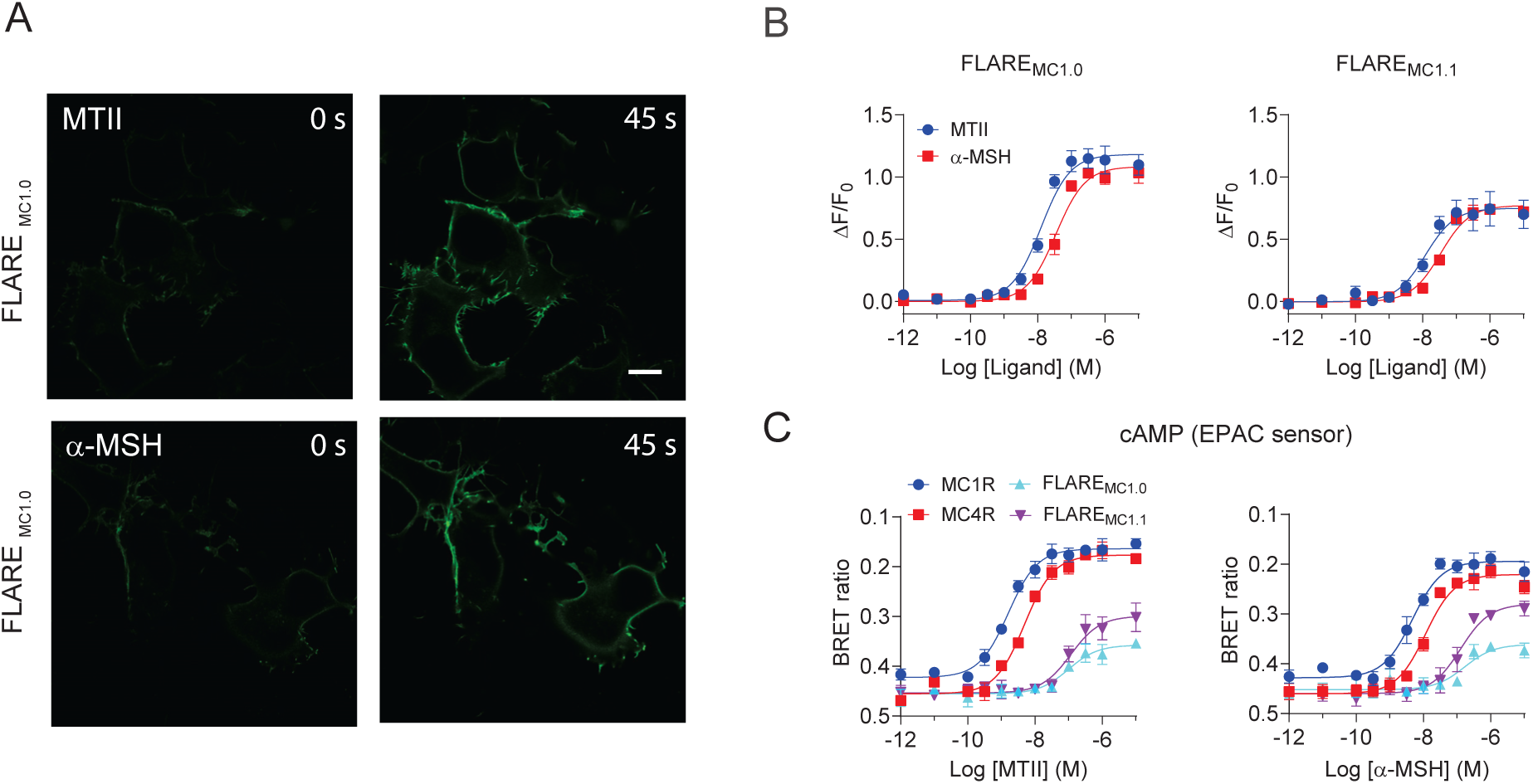
**Functional characterization of FLARE_MC_ sensors**. (**A**) Fluorescence increases upon receptor activation of FLARE_MC1.0_ expressed in HEK 293 cells. Confocal images of FLARE_MC1.0_ transfected cells before (0 s) and after 45 s stimulation with either 1 μM of MTII (*top*) or 2 μM of α-MSH (*bottom*). Scale bar, 10 μm. (**B**) Comparison of FLARE_MC1.0_ and FLARE_MC1.1_ responses. Cells expressing the FLARE_MC_ sensors in a 96 well plate were stimulated with various concentrations of either MTII (blue circle) or α-MSH (red square) for approximately 2 min, before fluorescence was measured. Fluorescence changes were expressed as βF/F_0_ and data are means ± SEM from 3 independent experiments. (**C**) cAMP responses from melanocortin receptors. HEK293 cells either expressing MC1R (blue circle), MC4R (red square), FLARE_MC1.0_ (cyan triangle), or FLARE_MC1.1_ (purple inverted-triangle) sensors along with EPAC BRET sensor, were stimulated with various concentrations of MTII (*left*) or α-MSH (*right*) for 3 min before signals were measured. Data are means ± SEM from 3 independent experiments.

### FLARE_MC_ is a selective and sensitive sensor for selectively detecting melanocortin peptides

We next investigated the selectivity of FLARE_MC_ sensor by assessing its responsiveness to other melanocortin peptides and to the natural inverse agonist AgRP. Beta-MSH activated the sensor with a potency comparable to that of a-MSH, while both ACTH and y_1_-MSH induced activation of FLARE_MC_, albeit with much lower potency (Figure 3A). These findings are consistent with previously reported relative affinities of these melanocortin peptides for MC1R (3). The mMC1R has reportedly lower affinity for the AgRP compared to mMC4R (3). Consistently, employing a competition assay, we found AgRP to be 75-100-fold more potent at inhibiting a-MSH-mediated signaling at MC4R, than at MC1R (Figure 3B). FLARE_MC_ showed similar insensitivity to AgRP (Figure 3C). Consistent with its higher specificity as an inverse agonist for MC4R vs MC1R, AgRP showed low efficacy and potency at reducing basal FLARE_MC_ fluorescence (Figure 3C, right panel). We evaluated a panel of reported MC1R ligands, including SHU9119, HS024, and MSG606, with varying selectivity and agonistic activity toward this receptor (24–27), as well as other neurotransmitters and peptides, to assess their effects on FLARE_MC_’s responses (Figure 3D). SHU9119, HS024, and MSG606 all elicited robust activation of the sensor, comparable to MTII. In contrast, a range of neurotransmitters and unrelated peptides, including dopamine, serotonin (5-HT), neuropeptide Y (NPY), angiotensin II, arginine vasopressin, oxytocin, and bradykinin, failed to activate the FLARE_MC_. These results support the specificity and sensitivity of the FLARE_MC_ for melanocortin peptides and synthetic ligands.

**Figure 3:**
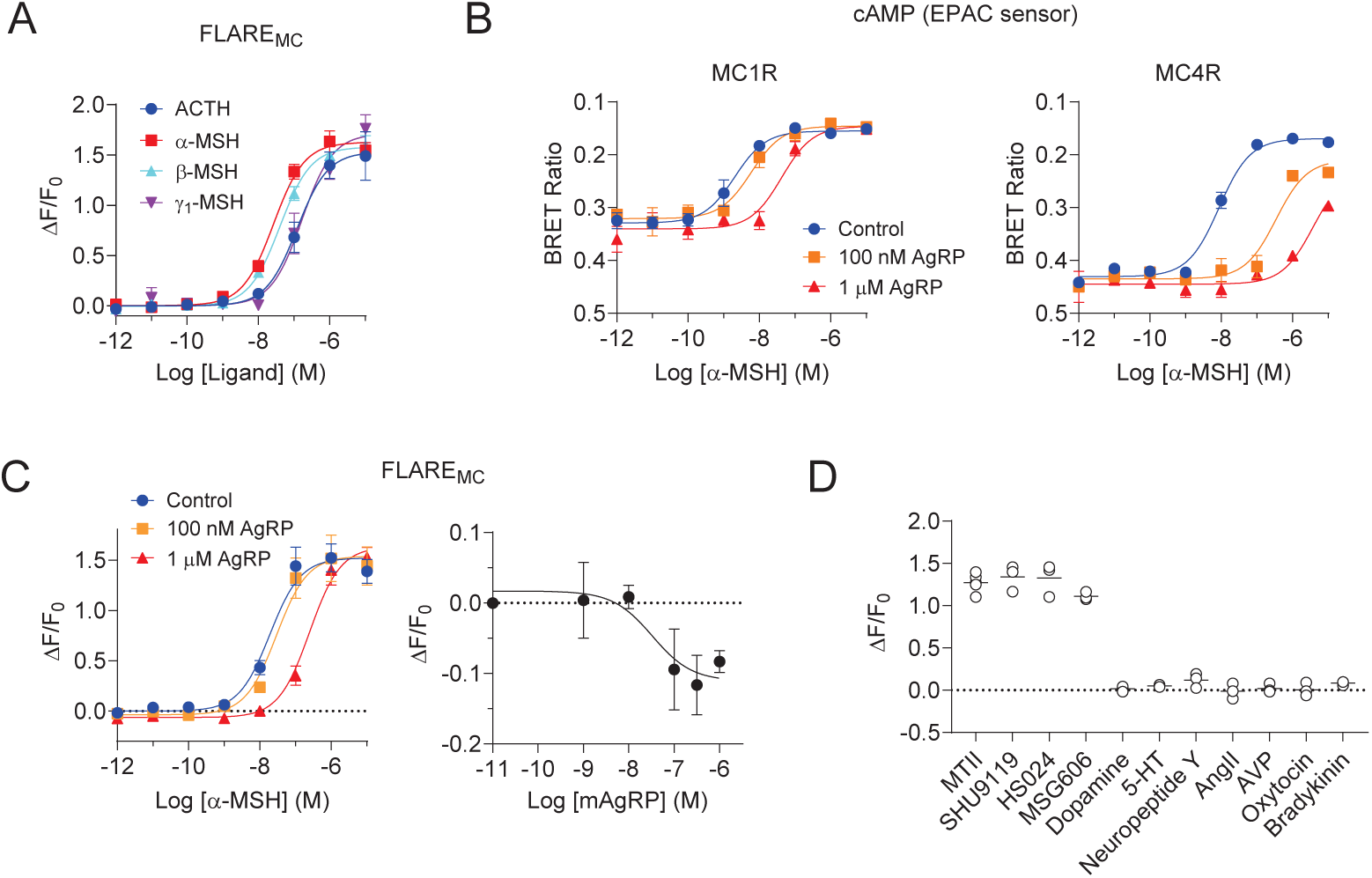
**Pharmacological characterization of FLARE_MC_ sensor**. (**A**) Endogenous melanocortin peptides increase FLARE_MC_ fluorescence in HEK293 cells. Cells expressing the FLARE_MC_ sensors in a 96 well plate were stimulated with various concentrations of ACTH, α-, β-, or ψ_1_-MSH for approximately 2 min, before fluorescence was measured. Fluorescence changes were expressed as βF/F_0_ and data are means ± SEM from 3 independent experiments. (**B**) Competing effects of AgRP on melanocortin receptors signaling. HEK293 cells expressing EPAC BRET sensor along with either MC1R (*left*) or MC4R (*right*) were incubated for 15 min in the presence or absence (control) of 100 nM or 1 μM mAgRP, followed by stimulation with various concentrations of α-MSH for 3 min prior to BRET measurements. Data are means ± SEM from 3 independent experiments. (**C**) Effects of AgRP on the FLARE_MC_’s responses. FLARE_MC_ transfected cells were incubated without (control) or with 100 nM or 1 μM mAgRP for 15 min and then stimulated with various concentrations of α-MSH for 3 min before fluorescence measurements (*left*). Cells expressing FLARE_MC_ were treated with various concentrations of mAgRP, before fluorescence measurements (*right*). Data are means ± SEM from 3 independent experiments. (**D**) Effects of different ligands on FLARE_MC_ responses. FLARE_MC_-expressing HEK293 cells in 96 well plates were stimulated with MTII (1 μM), SHU9119 (1 μM), Dopamine (100 μM), and HS024, MSG606, 5-HT, Neuropeptide Y, Angiotensin II (AngII), Arginine Vasopressin (AVP), Oxytocin, and Bradykinin (10 μM each). Data are means ± SEM from 3 independent experiments.

### FLARE_MC_ sensor does not recruit β-arrestin nor internalize

We tested the ability of the FLARE_MC_ to recruit β-arrestin and internalize which would limit its sensitivity in cells and *in vivo*. While MTII treatment resulted in MC1R-EGFP internalization, the FLARE_MC_ failed to internalize, and the sensor fluorescence remained detectable only at the cell surface (Figure 4A). We used the sensitive B-arrestin ebBRET assay, consisting of B-arrestin2-RlucII and rGFP-caax, to assess the extent of B-arrestin recruitment to FLARE_MC_ (28) (Figure 4B). Both MTII and α-MSH potently promoted B-arrestin recruitment to MC4R. Interestingly, α-MSH was more efficacious than MTII at promoting B-arrestin recruitment to MC4R. We observed no significant agonist-mediated B-arrestin recruitment to the MC1R, although a small consistent increase in signal was observed at high MTII concentrations. Beta-arrestin has been reported to bind to MC1R mostly without stimulation, and melanocortin only slightly increases this interaction after long term agonist stimulation (29). However, we found that FLARE_MC_ did not recruit B-arrestin in response to either MTII or a-MSH. Its lower basal B-arrestin recruitment, compared to MC1R and MC4R, may also reflect the sensor’s reduced affinity to B-arrestin.

**Figure 4:**
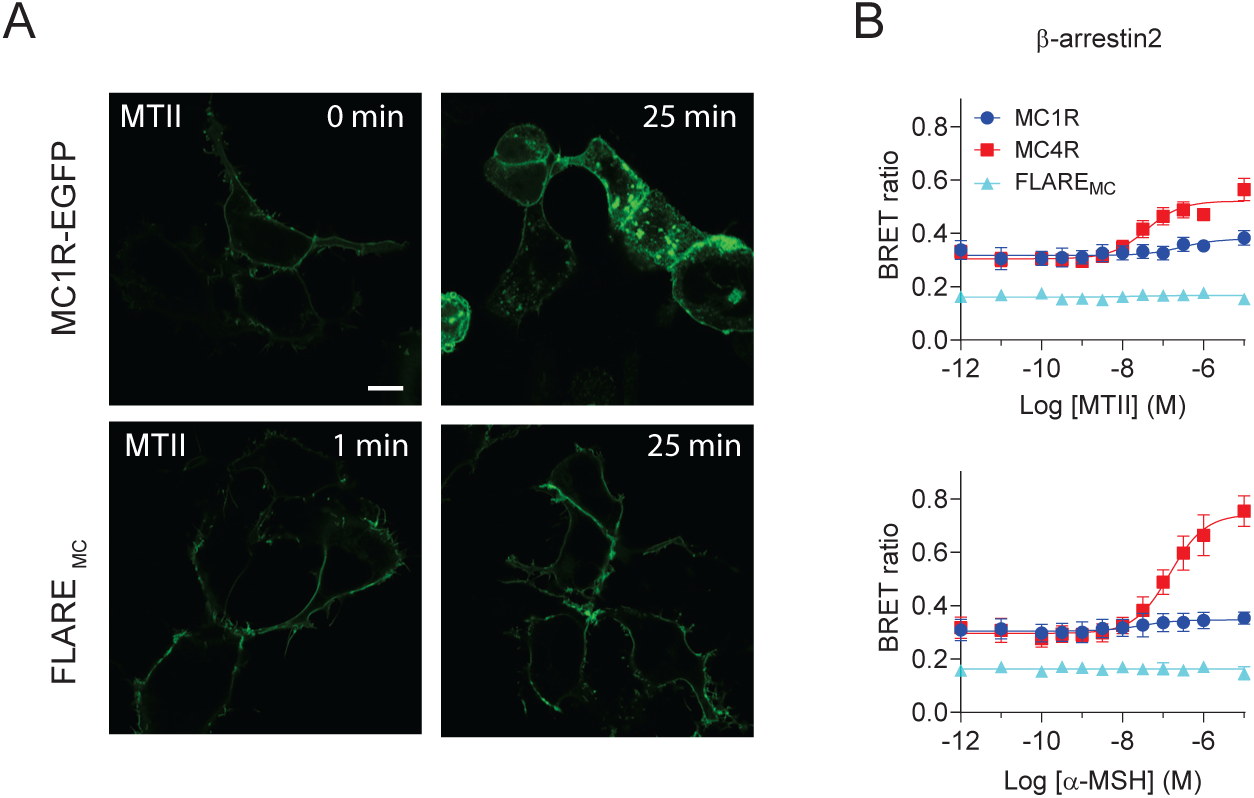
Assessment of β-arrestin interaction with MC1R, MC4R and FLARE_MC_, and internalization of receptors. (**A**) HEK293 cells were transfected either MC1R-EGFP or FLARE_MC_ sensor. Images were taken before (0 min), 1 min, and 25 min after incubation with 1 μM MTII at 37°C. Scale bar, 10 μm. (**B**) β-arrestin2 recruitment by melanocortin receptors. Cells transfected with MC1R (blue circle), MC4R (red square) or FLARE_MC_ (cyan triangle) along with the BRET pair β-arrestin2-RlucII and rGFP-caax were stimulated with various concentrations of either MTII (top) or α-MSH (bottom) for 5 min, before measurements. Data are means ± SEM from 3 independent experiments.

### The FLARE_MC_ sensors exhibits high photostability, efficient spectral separation, and reversible photoactivation

We next characterized key spectral properties of the sensor, including its excitation and emission spectra, photostability and kinetic profiles, important parameters to consider for longitudinal applications both in cells and *in vivo* studies. The one-photon spectral properties of FLARE_MC_, assessed at basal state (saline) and following MTII or a-MSH stimulation, revealed a peak excitation at 500 nm with an isosbestic point between 405-425 nm, and an emission maximum at 515 nm (Figure 5A), enabling efficient signal separation, comparable to other GRAB sensors. (12, 30, 31). To assess the photostability of the sensor, we continuously photoactivated FLARE_MC_-expressing HEK293 cells using one-photon excitation and monitored the fluorescence decay over time (Figure 5B). The FLARE_MC_ exhibited better stability than the EGFP-tagged MC1R and the GRAB_DA2m_ sensor. Despite a slight reduction in photostability of MTII-bound FLARE_MC_ compared to the unliganded sensor, MTII-FLARE_MC_ demonstrated greater photo-resilience than GRAB_DA2m_, whether bound or unbound to Quinpirole. To characterise sensor kinetics, we used a perfusion system to apply agonist or medium to HEK293 cells expressing FLARE_MC_ and to remove ligands via mild acid washes (Figure 5C). Adding MTII to cells rapidly increased FLARE_MC_ fluorescence with r_on_ of around 1.7 s. Mild acid buffer washes efficiently dissociated the ligand from FLARE_MC_ and reduced the sensor’s fluorescence. Under these conditions, we observed a r_off_ of approximately 43 seconds. The r_on_ and r_off_ are in the range of observed values for other GRAB sensor (11, 16, 32), considering the diffusion of the agonist in the buffer and its binding to the sensor, and its removal upon perfusion washes. FLARE_MC_ activation was fully reversible, and the sensor was robustly reactivated by MTII (Figure 5E). Notably, while acidic wash transiently reduced FLARE_MC_ fluorescence below baseline levels, fluorescence rapidly recovered upon return to physiological conditions, suggesting resiliency of the sensor to acidification.

**Figure 5:**
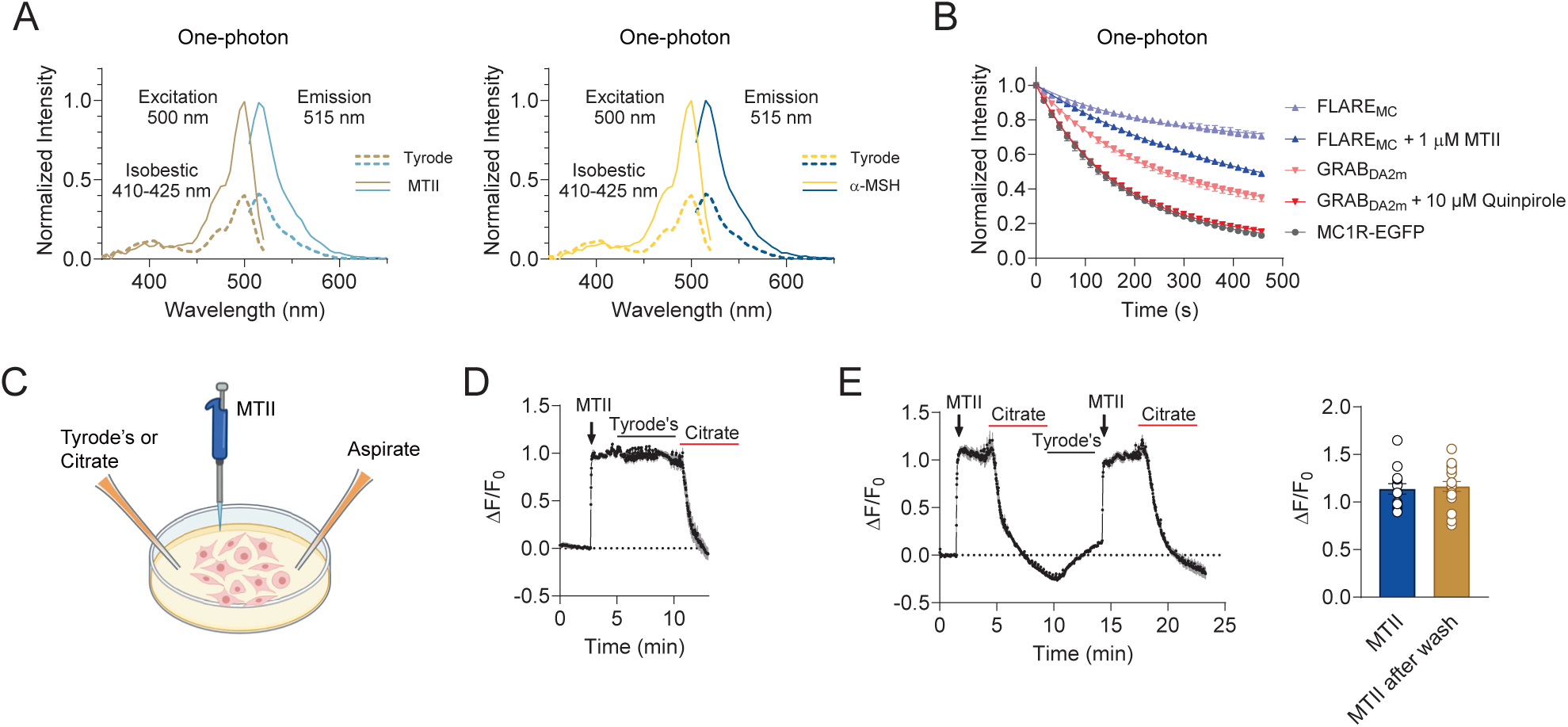
Photophysical properties of FLARE_MC_ sensor *in vitro*. (**A**) One-photon excitation and emission spectra of FLARE_MC_ sensor expressed in HEK293 cells in the absence (Tyrode’s buffer) or presence of 5 μM of MTII (*left*) or α-MSH (*right*). Each trace represents the mean of 4 independent experiments. Excitation and emission peaks are indicated. (**B**) One-photon photobleaching. Indicated labeled receptor or sensor DNA transfected cells were photobleached with or without corresponding ligands. Data are means ± SEM from 3 independent experiments. (**C**) Schematic diagram illustration of the perfusion systems. Tyrode’s buffer or citrate buffer (pH 5.2) were applied via gravity and the aspiration were done through a peristaltic pump. Image was created with www.biorender.com. (**D**) Fluorescence changes in FLARE_MC_ in response to 1 μM MTII application, followed by Tyrode’s buffer washing and then citrate buffer (pH 5.2) washing. Representative data of 2 experiments are shown, and the values are means signals of 3 cells from the same dish. (**E**) Reversibility of FLARE_MC_ fluorescence. Fluorescence changes in FLARE_MC_ in response to 1 μM MTII stimulation, with cycles of citrate, Tyrode’s washes and MTII stimulations. Representative traces from 3 independent experiments are shown (*left*), and the values are means of 3 cells in the same dish. Represented are the maximum fluorescence (βF/F_0_; *right*) of cells treated with MTII or acid washed and re-challenged with MTII. Data are from 3 independent experiments.

### Neuronally expressed FLARE_MC_ efficiently detects melanocortin in vivo and in vitro

We next examined whether the sensor maintained its sensitivity in a homologous expression system. To this end, we delivered a viral construct of FLARE_MC_ under a human synapsin promoter (hSYN; AAV-hSYN-FLARE_MC_), to the mouse neural crest-derived Neuro2a cell line, a well-established model for neuronal differentiation, neurite outgrowth, and intracellular signaling (33), and primary neurons (Figure 6A). FLARE_MC_ efficiently and rapidly detected melanocortin stimulation in both cell types. In Neuro2a cells, the sensor displayed potency and efficacy toward a-MSH and MTII comparable to those observed in HEK293 cells (Figure 6B and 2B), indicating that its sensitivity is largely dictated by intrinsic properties rather than the cellular environment.

**Figure 6:**
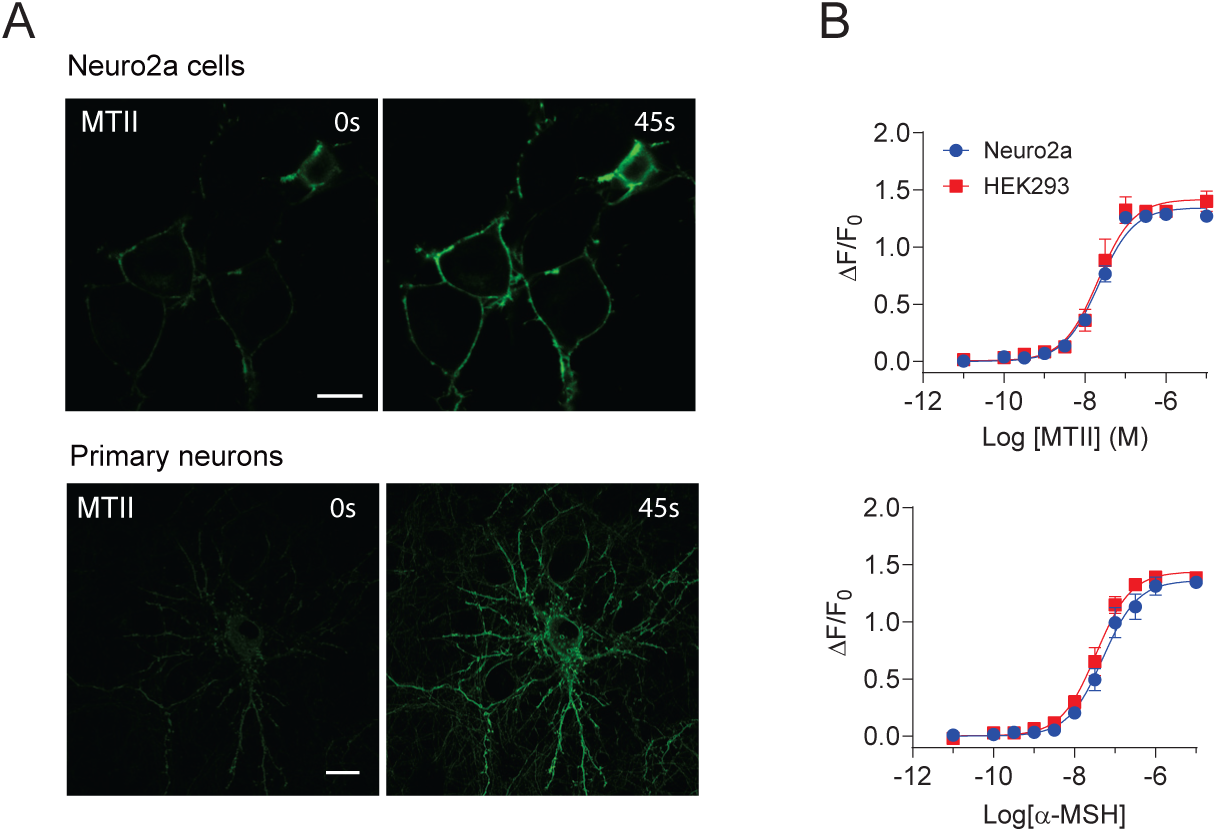
**Activity of FLARE_MC_ in neuronal cells**: (**A**) Confocal microscopy images of pAAV-hSyn-FLARE_MC_ DNA transfected Neuro2a cells (*top*) and AAV-hSyn-FLARE_MC_ transduced mouse primary cortical cells (*bottom*). Images were taken before (0 s) and 45 s after MTII (final concentration of 5 μM) stimulation. Scale bar, 10 μm. (**B**) Concentration response curves of FLARE_MC_ for MTII (*top*) and α-MSH (*bottom*) in Neuro2a cells and HEK293 cells. Cells expressing the FLARE_MC_ sensors in a 96 well plate were stimulated with various concentrations of either MTII (blue circle) or α-MSH (red square) for approximately 2 min, before fluorescence was measured. Fluorescence changes were expressed as ΔF/F_0_ and data are means ± SEM from 3 independent experiments.

Finally, we determined the extent to which FLARE_MC_ could reliably detect changes in melanocortin signaling in relevant CNS sites for melanocortin signaling. To this end, we delivered the packaged AAV-hSYN-FLARE_MC_ into the PVN of wildtype mice, which led to robust expression of the melanocortin sensor (Figure 7A, B). To assess ligand-induced fluorescence changes of the sensor *in vivo*, optic fibers were implanted above the PVN at the time of virus injection (Figure 7C). To determine if FLARE_MC_ could detect pharmacological increases in melanocortin, we systemically administered MTII to these animals during the fiberphotometric recordings. As compared to saline injected animals, MTII significantly increased FLARE_MC_ fluorescence immediately after injection with peak fluorescence observed at 30 minutes post drug delivery, and the signal persisting for an hour post injection (Figure 7D, E), suggesting that the FLARE_MC_ is a suitable tool for studying melanocortin release dynamics *in vivo*.

**Figure 7:**
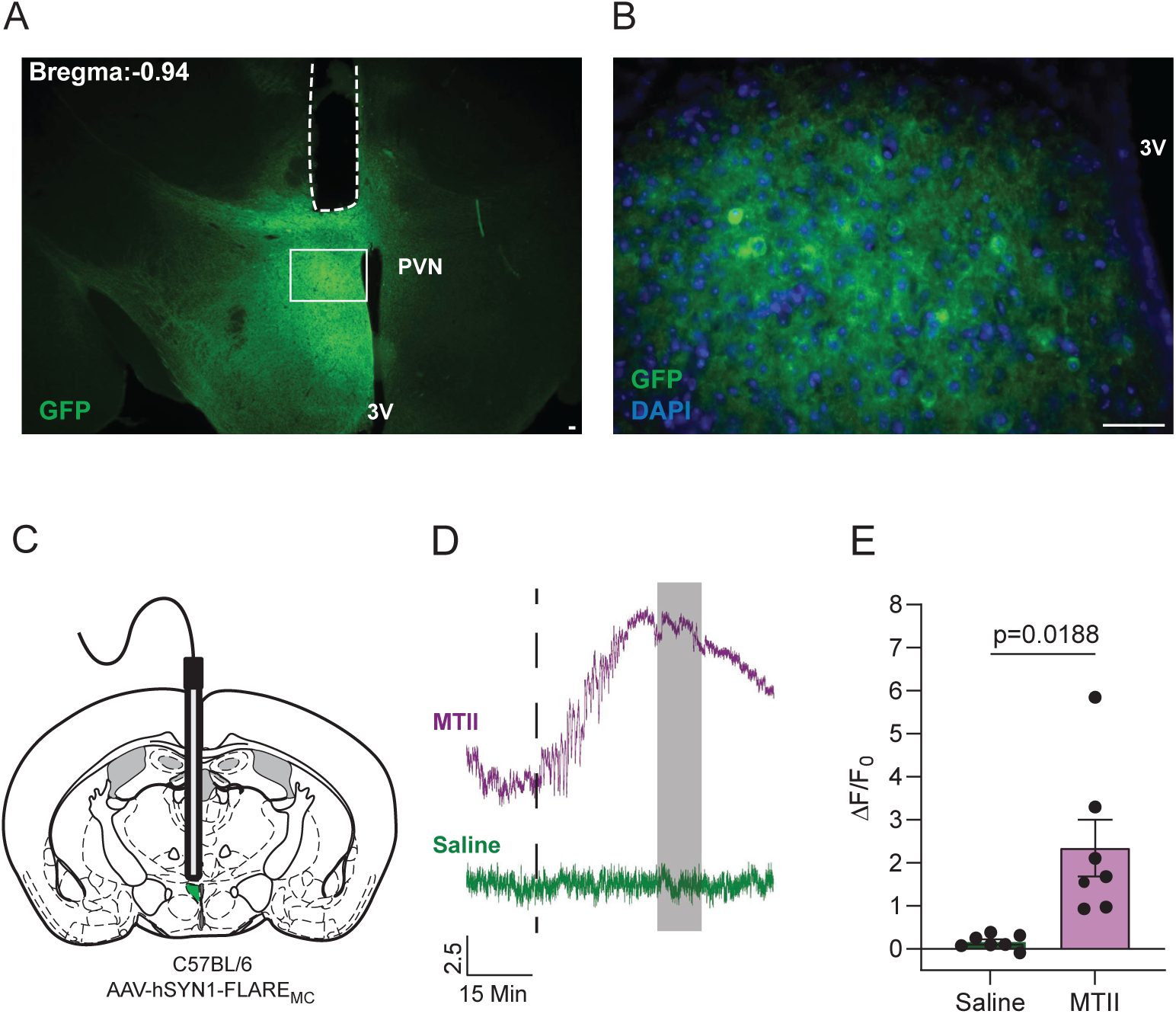
Intravital imaging of FLARE_MC_ in the hypothalamus. (**A**) Representative image of GFP immunostaining (green) of a hypothalamic section from a wildtype C57BL/6 mouse transduced with AAV-hSYN-FLARE_MC_ in the PVN region. Position of the recording fiber implant is indicated by dashed line. (**B**) Higher resolution display of the area boxed in **A**, showing immunoreactive FLARE_MC_ (green) and DAPI-stained cell nuclei (blue). Scale bars (**A** and **B**), 20 μm. (**C**) Schematic depicting the recording site in mouse brain. Shown is the optic fiber implant above the PVN transduced with the sensor-expressing virus (green). (**D**) Representative recording of FLARE_MC_ activity in response to intraperitoneal injection (dashed line) of either saline or MTII (8 µg/g body weight), and (**E**) quantification of fluorescence signals (ΔF/F_0_) at 30-40 min (gray bar in **D**) post treatments after baseline subtraction. PVN, paraventricular nucleus of the hypothalamus; 3V, third ventricle. Data are presented as mean ± SEM and analyzed with a paired t-test; n=7.

**Figure 8:**
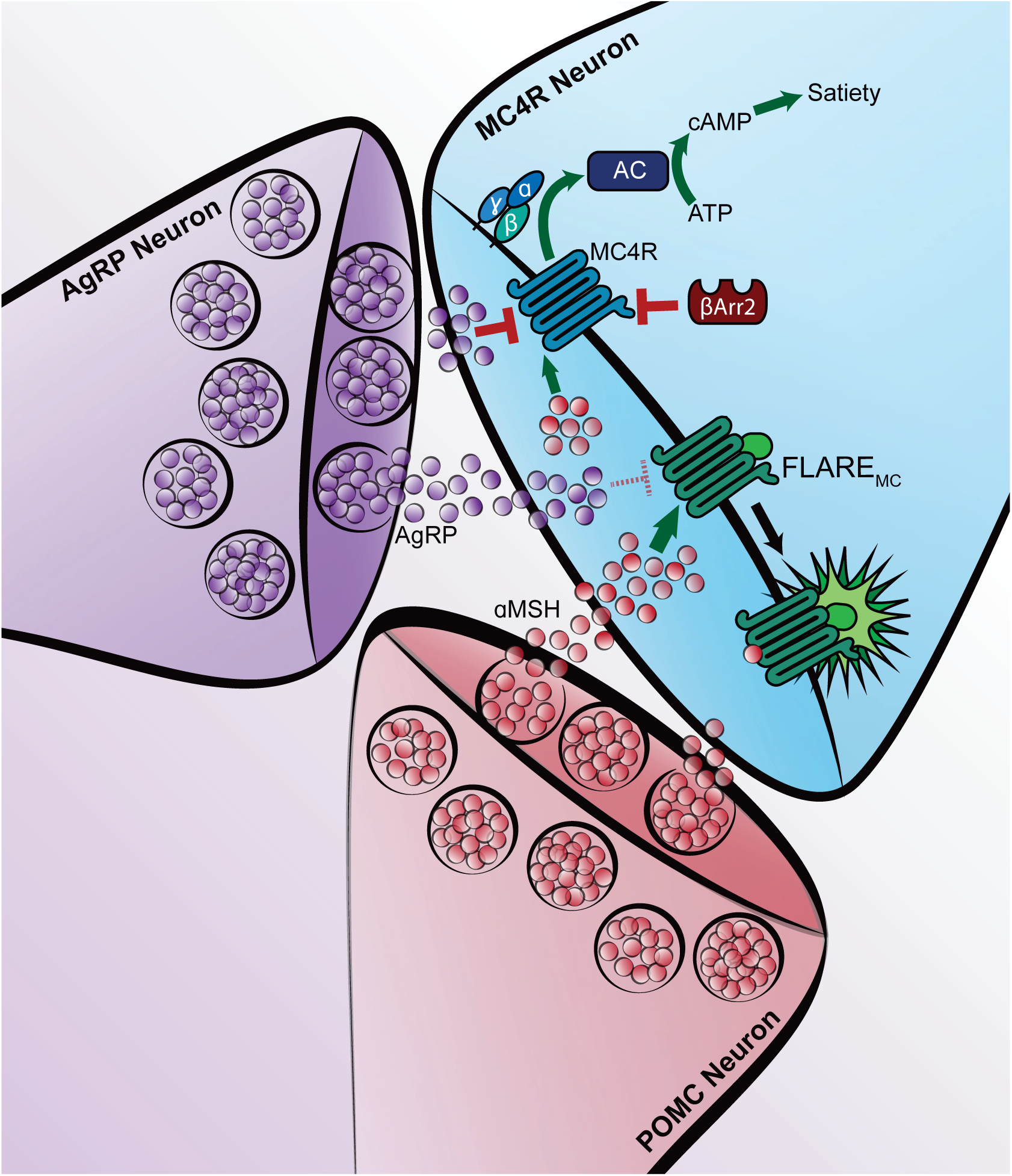
**Melanocortin peptide sensing in the brain using FLARE_MC_**. The PVN receives input from both AgRP and POMC neurons which signal through the Gs-coupled MC4R. POMC derived peptides, including α-MSH, activate MC4R resulting in cAMP production via Adenylate cyclase (AC) and satiation, whereas the inverse agonist AgRP reduce MC4R-based signaling resulting in hunger. MC4R signaling is desensitized by B-Arrestin2 (BArr2). The FLARE_MC_ sensor, derived from MC1R, binds POMC peptides with higher affinity relative to AgRP but does not result in intracellular signaling, BArr2 desensitization or internalization.

## Discussion

Here we report the development and characterization of FLARE_MC_, a highly selective, and sensitive genetically-encoded sensor for the melanocortin a-MSH, which is key to energy balance control. This sensor, the first of its kind for detecting a peptide controlling appetite, is designed for real-time *in vivo* detection of endogenous melanocortin release. Adding to the tools available for monitoring neurotransmitter regulation (34), the FLARE_MC_ sensor will unlock new insights into the role of central melanocortin signaling in appetite and body weight regulation. FLARE_MC_ offers several distinct benefits. First, it exhibits high sensitivity and selectivity for melanocortin peptides in the nanomolar range, but not other peptides or neurotransmitters, with a potency rank *in vitro* of a-MSH > B-MSH >> y-MSH = ACTH, consistent with MC1R’s high affinity for melanocortin and analogues (3). Second, unlike the use of a MC4R-based sensor, the MC1R-based FLARE_MC_ is less responsive to AgRP. Given the rank order of potency and the fact that rodents do not produce B-MSH in the hypothalamus, the sensor will most likely detect a-MSH dynamics in the PVN (35). Third, the sensor binding is reversible, demonstrated by fast on and off kinetics, in support of effective real-time monitoring of melanocortin actions over extended periods. Forth, melanocortin binding to FLARE_MC_ sensor at physiological levels will likely produce minimal downstream G protein signaling, hence ensuring negligible interference with the natural MC4R tone in the hypothalamus and the detection of the anorexigenic a-MSH release in the brain. Although we have not specifically investigated the mechanism of desensitization for FLARE_MC_, our findings indicate that the sensor does not internalize and exhibits minimal downstream signaling. These properties suggest the FLARE_MC_ is highly suitable for long-term *in vivo* studies without signal loss due to either sensor desensitization or downregulation. Future work should be directed at designing FLARE_MC_ variants capable of detecting both AgRP and a-MSH responses, or AgRP selectively. Such sensors, in conjunction with the specific melanocortin sensor developed here, would enable a more comprehensive investigation of the *in vivo* interplay between a-MSH and AgRP in regulating appetite.

We further demonstrated the suitability of the sensor for *in vivo* studies. Photometric quantification of FLARE_MC_ expressed in the PVN revealed an increase in fluorescence upon peripheral delivery of the synthetic a-MSH analog MTII. This finding indicates that the sensor is able to detect circulating ligands when expressed in neurons of the PVN region, which is the main target of arcuate nucleus POMC neurons to convey their anorexigenic drive (36, 37). Although the FLARE_MC_ showed similar pharmacological responses to melanocortin peptides in various cell types, including cultured neurons, its *in vivo* sensitivity may vary due to expression levels and cellular context (38). Nevertheless, our *in vivo* findings on sustained melanocortin-induced activation in the PVN and the photonic resilience of the sensor supports its usefulness for longitudinal studies. Therefore, future experiments could involve exploring the ability of the sensor to detect changes in extracellular levels of endogenous a-MSH across the daily cycle and the contributions of homeostatic versus non-homeostatic drives (39), regulating feeding and fat mass accumulation (40, 41). It could also provide critical insights into if, how, and when the melanocortin system fails to prevent diet-induced obesity (DIO). Additionally, this sensor could uncover the underlying mechanisms of leptin resistance in DIO, a research area of considerable clinical importance (42–45). This is specifically relevant considering that MC4R agonists are seemingly effective in treating genetic but not DIO obesity (41).

In summary, FLARE_MC_ serves as a valuable complement to sensors that target other feeding-related systems, such as dopamine sensors (46, 47) which probe the reward circuitry. In combination, these sensors could critically help in elucidating the relative contribution of the reward versus homeostatic systems to feeding and body weight control (48). Additionally, FLARE_MC_ should significantly enhance screening for effective melanocortin receptor ligands *in vivo*, furthering treatment strategies for obesity, anorexia, and psychiatric conditions (41, 49).

## Experimental Procedures

### Materials

Melanotan II (MTII) was purchased from Genscript (Piscataway, NJ, USA). α-MSH, β-MSH, ψ_1_-MSH, ACTH, MSG606, Neuropeptide Y were from Cayman chemicals (Ann Arbor, MI, USA). Mouse AgRP (82–131) and SHU9119 were obtained from Pheonix pharmaceuticals (Burlingame, CA, USA). HS024 and MTII were obtained from Bachem AG (Bubendorf, Switzerland). Dopamine, Angiotensin II (AngII), Arginine Vasopressin (AVP), Oxytocin, 5-HT, and other chemicals were obtained from Sigma-Aldrich (St. Louis, MO, USA). Dulbecco’s modified Eagles medium (DMEM), fetal bovine serum (FBS), and other cell culture reagents were purchased from Gibco (Thermo Fisher Scientific, Grand Island, NY, USA). Coelenterazine 400a was purchased from Nanolight Technology (Prolume Ltd., Pinetop, AZ, USA). Restriction enzymes, Q5 DNA polymerase, and Gibson assembly mix were obtained from New England Biolabs (NEB, Ipswich, MA, USA). Oligonucleotides were synthesized at Integrated DNA Technologies (Coralville, IA, USA). pAAV-hSyn-GRAB-DA2m (16) (addgene #140553), pDisplay-GRAB_eCB2.0-IRES-mCherry-CAAX (11) (addgene #164611), and pAAV-hSyn-GRAB-NE1m (12) (addgene #123308) were obtained from addgene (www.addgene.org, Watertown, MA, USA). Polyethylenimine (linear, MW 25,000) was purchased from Polysciences, inc. (Warrington, PA, USA).

### DNA constructs and mutagenesis

For general expressions in mammalian cells, GRAB_DA2m_ was subcloned from pAAV-hSyn-GRAB_DA2m_ into HindIII/XhoI sites of pcDNA3.1/zeo(+) by PCR amplification and Gibson assembly. Mouse MC1R and MC4R cDNA’s were obtained via PCR using mouse genomic DNA as a template and subcloned into pcDNA3.1/zeo(+) vector which containing Igk-leader sequences. MC1R-EGFP and MC4R-EGFP were generated by overlapping PCR of products of amplified MC1R or MC4R with PCR amplified from EGFP and Gibson assembly into HindIII/XhoI sites in pcDNA3.1/zeo(+). CpGFP with ICL3 linker of each GRAB sensors were PCR amplified by using GRAB_DA2m_, GRAB_eCB2.0_ and GRAB_NE1m_ sensor DNA as a template and subcloned into MC1R’s ICL3 by overlapping PCR and Gibson assembly. For sensor packaging into AAV, FLARE_MC_ in pcDNA3.1/zeo(+) was PCR amplified using AgeI-Igkleader-F and EcoRI-RV-HindIII-R primers and Gibson assembled into AgeI/EcoRI sites in pAAV-hSynGRAB_DA2m_ DNA. Mutagenesis was done by overlapping PCR with a NBB or NNB sequences in the primers. AAV-FLARE_MC_ was packaged in serotype 8 (Neurophotonics, Quebec, QC, Canada) at a titer of 6.2×10^12^ GC/ml. All primers used in this study are listed in supplementary table 1. DNA sequences were verified by Sanger sequencing (Genome Quebec, Montreal, QC, Canada).

### Cell culture and transfection

HEK293 and Neuro2a cells were cultured in DMEM supplemented with 10% FBS and 20 µg/ml gentamicin. Cells were grown at 37 °C in 5% CO_2_ and 90% humidity. Cells were seeded at a density of 150,000 cells/well in a 6 well plate (or glass-bottom 35 mm dish for imaging) or 9,000 cells/well in the poly-L-ornithine coated 96 well plate and next day transfected with PEI methods. Briefly, 1 µg of total DNA in 100 µl of PBS were mixed with 100 µl of PBS containing 2∼3 µl of 1 mg/ml of PEI. For fluorescence measurement, 60 ng FLARE_MC_ sensors DNA were used. For cAMP measurement with EPAC BRET biosensor, 100 ng of receptor DNA along with 50 ng of EPAC sensor (GFP10-EPAC-RlucII) were used. For β-arrestin2 translocation assay, 100 ng of receptor DNA along with 20 ng of β-arrestin2-RlucII and 80 ng of rGFP-caax were used. Total DNA amount was adjusted to 1 µg with empty vector DNA (pcDNA3.1/zeo(+)). After 20 min incubation, the DNA/PEI complexes were dispensed onto cells in 96 well plates (15 µl/well) or added (200 µl/well) onto cells in 6 well plate or 35 mm dish (for confocal microscopy). For the sensor variants screening, cells were seeded at a density of 9,000 cells/well in a poly-L-ornithine coated black 96 well plate and were transfected the next day. For the BRET assay, cells were seeded on a poly-L-ornithine coated white 96 well plate. Transfection of Neuro2a cells in a glass bottom 35 mm dish was done with 300 ng of pAAV-hSyn-FLARE_MC_. All assays were performed 48 h post-transfection.

### Fluorescence measurements

Cells in 6 well plates were dissociated with TrypLE, and resuspended in Tyrode’s buffer (140 mM NaCl, 2.7 mM KCl, 1 mM CaCl_2_, 12 mM NaHCO_3_, 5.6 mM D-glucose, 0.5 mM MgCl_2_, 0.37 mM NaH_2_PO_4_, 25 mM HEPES, pH 7.4) and distributed onto black 96 well plates (∼50,000-80,000 cells/well) or cells transfected in 96 wells were washed once with Tyrode’s buffer. Cells were stimulated without (Tyrode’s buffer, control) or with a single concentration (final concentration of 10 μM for the screening) or various concentrations of MTII or α-MSH and fluorescent intensity was measured using a microplate reader (Infinite M200 pro, Tecan) at an excitation of 486/9 nm and emission of 535/20 nm. To measure specific fluorescence (F) intensity, values from mock (empty pcDNA vector) transfected cells were subtracted from values of all the sensor transfected cells. F_0_ represents the specific F value in the absence of the ligand (control), while ΔF indicates the specific F value in the presence of the ligand, with the F_0_ value subtracted. Changes in fluorescence were quantified by expressing ΔF divided by F_0_ (ΔF/F_0_). One-photon excitation and emission spectra were measured in FLARE_MC_ expressing HEK293 cells with or without 5 μM of MTII or α-MSH using the Tecan M200 Pro plate reader. Excitation spectra were acquired from 350 to 520 nm with a 5 nm step size and a 10 nm bandwidth, while emission was set at 560 nm with a 20 nm bandwidth. Emission spectra were measured from 505 to 650 nm with a step size of 5 nm and a bandwidth of 20 nm, while excitation was set to 470 nm with a10 nm bandwidth. Control cells transfected with an empty vector were used for background subtraction. After subtracting the background values, the fluorescent values were normalized to the peak excitation (500 nm) and emission (515 nm) of the ligand bound state of the sensor.

### Confocal microscopy

Before imaging, cells cultured in a glass-bottom 35 mm dish were rinsed once with Tyrode’s buffer (140 mM NaCl, 2.7 mM KCl, 1 mM CaCl_2_, 12 mM NaHCO_3_, 5.6 mM D-glucose, 0.5 mM MgCl_2_, 0.37 mM NaH_2_PO_4_, 25 mM HEPES, pH 7.4) and left in 1 ml of Tyrode’s buffer. Images of live cells were obtained on a Zeiss LSM-510 Meta laser scanning microscope with a Plan-Apochromat 63x/1.4 oil DIC objective lens using excitation/emission filter sets: 488 nm/ 505 nm (long pass). Ligands were applied in bolus to the cells using a micropipette, precisely achieving the intended final concentrations. To measure photostability, the cells were photobleached by illuminating a 488-nm laser with laser power of ∼290 µW and pixel dwell time of 12.8 µs. Cells were imaged with Plan-Neofluar 40x/1.3 Oil DIC objective lens. Mean ROI were analyzed by using Zen 2009 software. For the perfusion experiments, HEK293 cells were transfected in 6 well plates and following day, dissociated and re-plate onto poly-L-ornithine coated 42 mm glass coverslips in 60 mm dishes. After another day of incubation, the coverslips were mounted onto a perfusion chamber (POC-R2 cell Cultivation System, PeCon GmbH, Ziegeleistraße, Germany) with 2 ml of Tyrode’s buffer. For ligand wash off, citrate buffer (50 mM sodium citrate, 90 mM NaCl, pH 5.2) was used. The images were obtained with Plan-Apochromat 63x/1.40 oil DIC M27 objective lens on LSM 780, AxioObserver. Mean ROI were analyzed using Zen 2012 software.

### BRET assay

HEK cells transfected in a white 96 well plate were washed once with Tyrode’s buffer and left in Tyrode’s buffer. Coelenterazine 400a (final concentrations of 2.5 μM) was added 1 min before ligand stimulation. Cells were then stimulated with various concentrations of ligand for 3 min for the EPAC sensor or 5 min for the β-arrestin translocation sensor, prior to BRET measurements. BRET signals were measured using a Synergy2 (BioTek) microplate reader with emission filters, 410/80 nm and 515/30 nm for detecting the RlucII (*Renilla* luciferase) (donor) and GFP10/rGFP (acceptor) light emissions, respectively. The BRET ratio was determined by calculating the ratio of the light emitted by GFP10/rGFP over the light emitted by the RlucII.

### Primary mouse cortical neuronal culture

Cortical neurons were prepared from embryos (E15-16) of pregnant C57BL/6 mice as previously described (50) with some modifications. Briefly, embryonic brains were kept in hibernation media (Hibernate-E, Invitrogen) supplemented with 2% NeuroCult™ SM1 (Stemcell technologies inc, Vancouver, BC, Canada) and 0.5 mM L-GlutaMAX (Gibco) at 4 °C. Cortices were then micro-dissected in ice-cold HBSS (Gibco, 14170-112) supplemented with 33 mM sucrose, 10 mM HEPES, and 0.5 mg/ml penicillin/streptomycin/glutamine (Gibco, 10378-16). Tissue was trypsinized and triturated using P1000 pipette in supplemented neurobasal medium (2% NeuroCult™ SM1 and 0.5 mM L-GlutaMAX). Cells were seeded at a density of 500,000/dish on poly-D-lysine-precoated glass bottom 35 mm dishes. Neurons were infected with AAV-FLARE_MC_ at 10,000-60,000 vg per cell after 8 days culture *in vitro* (DiV). Confocal microscopy was performed 3 and 5 days after AAV infection.

### Mice

C57BL/6 (Jackson Laboratories, Bar Harbor, ME, USA) mice were housed at room temperature under a 12-hour light/12-hour dark cycle (lights on at 7 am) and received daily health checks. Mice had ad libitum access to water and regular chow (2018s, Harlan Laboratories, Indianapolis, IN, USA). All animal procedures were carried out in accordance with the recommendations of the Canadian Council on Animal Care (CCAC) and have been approved by the local McGill University Animal Care Committee.

### Surgery

Mice, age 10-12 weeks old, were anesthetized with isoflurane, and placed into a stereotax frame (Kopf Instruments, Tujunga, CA, USA). Two hundred nl of AAV-hSYN-FLARE_MC_ was injected unilaterally into the PVN (-0.8 mm A/P, 0.25 mm M/L, and-4.8 mm D/V) via glass capillary using a Nanoject III (Drummond, Broomall, PA, USA) at a flow rate of 10 nl/s. Immediately following AAV injection, a fiber optic probe (Doric Lenses, Quebec, QC, Canada) was implanted 100 μm above the viral injection coordinates and secured in place using Metabond (Parkell, <u>Edgewood, NY, USA</u>) and dental cement. To allow sufficient time for viral expression, all fiber photometry studies were performed at least four weeks post-surgery.

### Immunohistochemistry

Animals were euthanized with isoflurane anesthesia followed by CO_2_ asphyxiation and then transcardially perfused with 1x PBS for 5 min followed by 10% formalin for an additional 10 min. Brains were then collected and post-fixed in 10% formalin for 24 hours followed by 24 hours in 30% sucrose. Brains were then sectioned on a microtome as 30 µm thick free-floating sections. Sections were washed 3 times with PBS, blocked in 3% donkey serum in PBS supplemented with 0.1% Triton-X (PBST) for 2 hours. After blocking, sections were incubated in primary antibody (Aves anti-GFP cat#GFP-1020; Thermo Fisher, Waltham, MA, USA) in PBST for 24 hours at room temperature. The following day, tissues were washed three times with PBS and incubated for two hours with secondary antibody (goat anti-chicken FITC; Jackson Immunoresearch, Baltimore Pike, PA, USA) with DAPI (Tocris Bioscience, Bristol, UK). Tissues were then washed three times, mounted onto glass slides, mounting media and coverslips applied and imaged on an Olympus BX61.

### Fiber photometry and MTII treatment

A 2-color fiber photometry system was used (Doric Lenses, Quebec, Canada) to probe sensor responses *in vivo*. Photometry experiments were performed in sated mice. For intraperitoneal injections, MTII (Bachem, Bubendorf, Switzerland) was dissolved in saline and given at a dose of 8 µg/g body weight. MTII-dependent fluorescence changes were recorded using a 470 nm excitation light source (0.4-1 V at 208 Hz) while excitation at 405 nm (0.4-0.8 V at 572 Hz), the isobestic point, produced a ligand insensitive control signal to account for hemodynamic changes and ligand-independent recording artifacts such as animal motion (51, 52). Fluorescence was detected using a single photodetector (Model 2151 Newport Visible Femtowatt, Doric Lenses, Quebec, Canada). The recorded signal was demodulated and saved using a decimation factor of 100 employing Doric Neuroscience Studio (version 6). Recordings were conducted according to an automated non-continuous 1 min on/off regimen.

### Data analysis

Fluorescence changes of the sensors (ΔF/F_0_) and BRET curves were fitted using GraphPad Prim 10 software (GraphPad Software Inc., Boston, MA, USA). For *in vivo* data analysis, custom Python scripts were used. The control 405 nm isosbestic signal was aligned to the 470 nm signal using a least-squares linear fit. The ΔF/F_0_ was obtained using the formula “(470 nm – fitted 405 nm) / fitted 405 nm“. Changes in fluorescence (control and treatment) were calculated by subtracting the average ΔF/F value between-10 and 0 min prior injection from the value between 30 and 40 min post injection.

## Acknowledgments

We would like to thank Dr. Lenka Schorova, Ms. Nathalie Bedard and Dr. Simon Wing for help in preparing the primary cortical neurons.

## Credit authorship contribution statement

**Yoon Namkung**: Writing – original draft, Writing – review and editing, Formal analysis, Methodology, Conceptualization, Data curation. **Tal Slutzki:** Methodology, Writing – original draft, Formal analysis. **Joao Pedroso**, Methodology, Formal analysis. **Xiaohong Liu**, Methodology. **Paul Sabatini:** Methodology, Writing – original draft, Writing – review and editing, Formal analysis, Summary Figure Generation, Conceptualization, Funding acquisition. **Maia V. Kokoeva**: Writing – original draft, Writing – review and editing, Formal analysis, Conceptualization, Funding acquisition. **Stéphane A. Laporte:** Writing – original draft, Writing – review and editing, Formal analysis, Conceptualization, Funding acquisition.

## Supporting information

**Figure S1:**
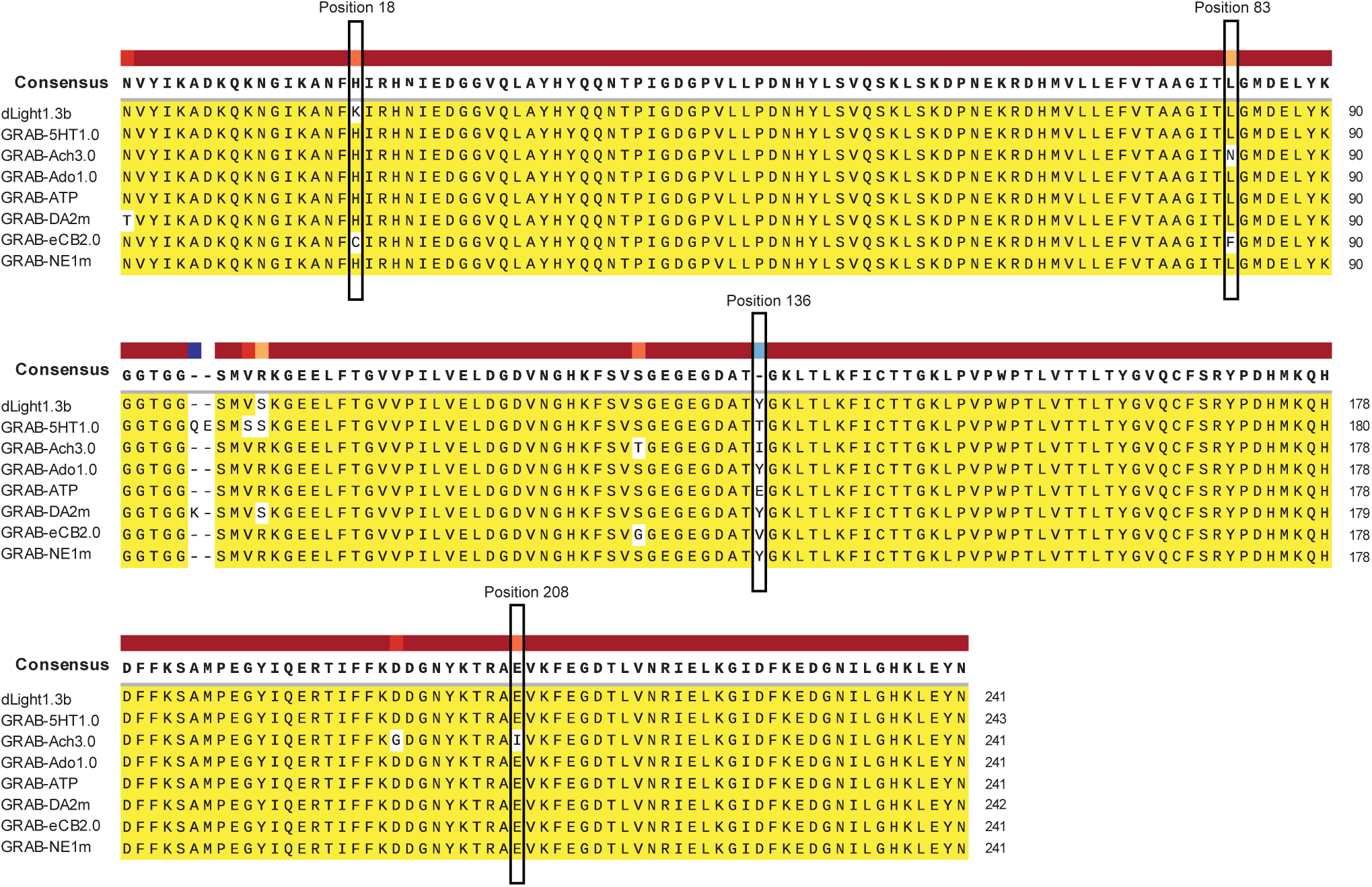
CpGFP amino acid alignment of various sensors. Targeted positions for the saturation mutagenesis are depicted in boxes.

**Figure S2:**
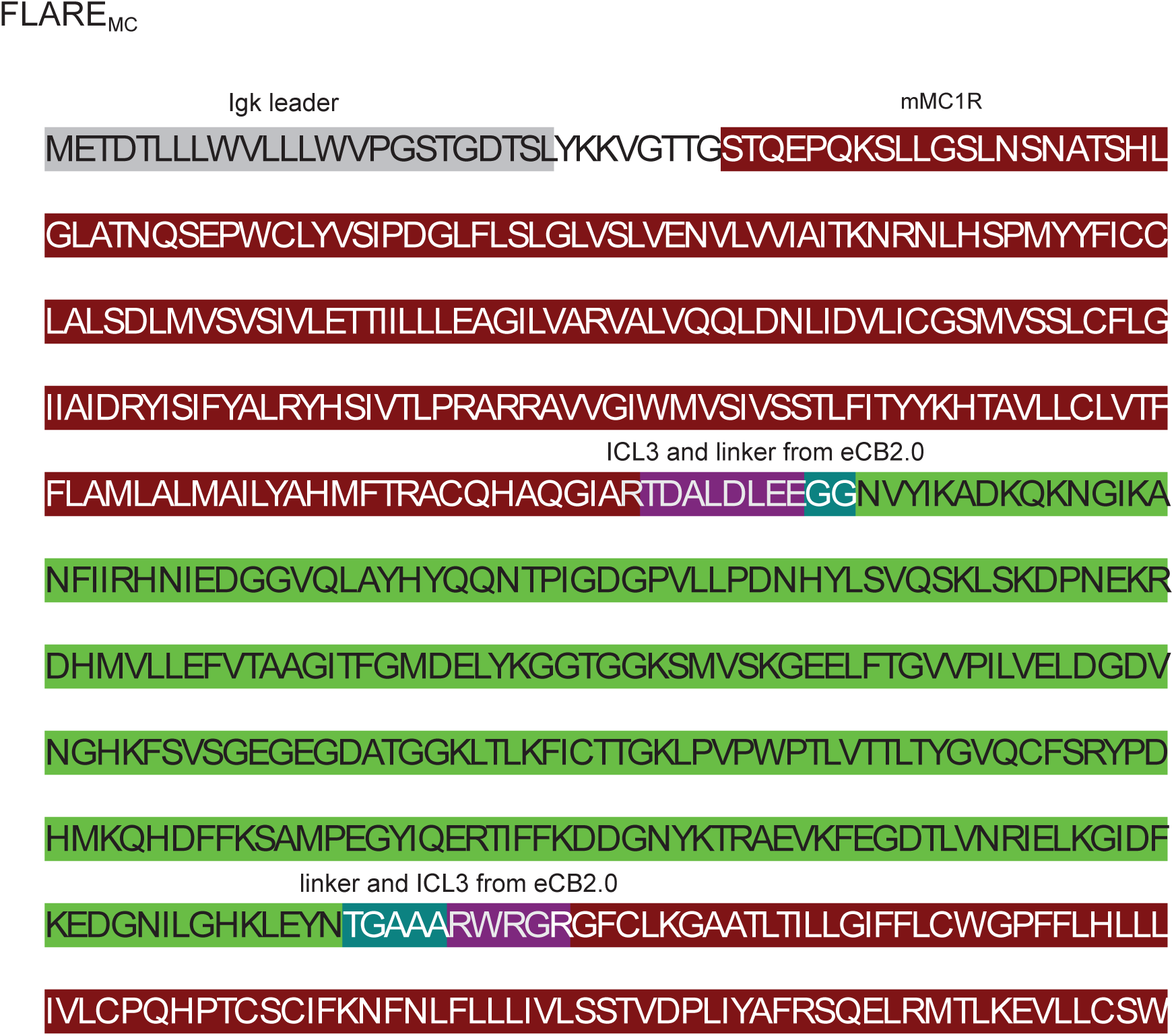
**Amino acid sequences of the FLARE_MC_ construct.**

---

**Supplementary Table 1.**
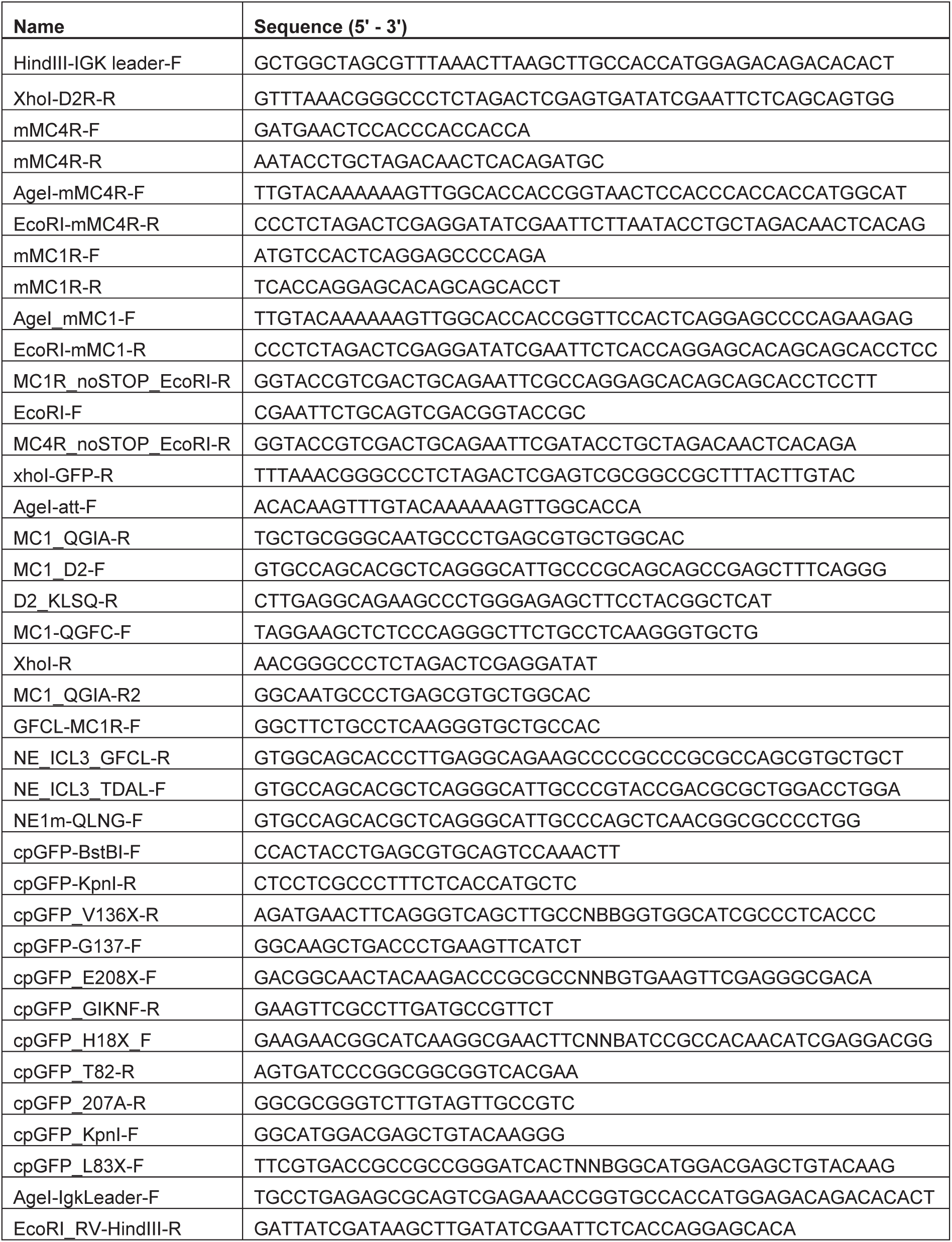
Primer sequences.

